# RePurpose: An LLM-Assisted, Source-Traceable Drug Repurposing Tool with Retrospective Validation on Clemastine in Multiple Sclerosis

**DOI:** 10.64898/2026.10.14.759218

**Authors:** Nitin Nayagam, Akanksha Arun

**Affiliations:** Rowan-Cabarrus Community College, Concord, NC 28027, USA; Sirpi Products and Services Private Limited, Bangalore, 560024, India

## Abstract

**Summary:** Thus far, evidence for drug repurposing is contained in target-, drug-, trial-, and literature-based data sources that are challenging to synthesize for non-specialists. Here, we introduce RePurpose, a web-based tool that takes as input a topic written in plain language and returns candidate drugs built from data in Open Targets, ClinicalTrials.gov, UniProt, and PubMed. For queries in the form of a goal, RePurpose filters candidate drugs by whether they modulate all targets implicated in the goal in the corresponding direction. Every candidate has a source that contributes evidence: a structured database, a clinical trial, or a proposal from a large language model. An adjustable ranking adjusts the order of candidates from well-studied to less well-known candidate drugs. We retrospectively tested RePurpose with the trial drug clemastine, which was being considered as a remyelinating agent in multiple sclerosis (MS). RePurpose retrieved clemastine in each replicate of the mechanism-worded queries, but in no replicates of the disease-worded queries, and no structured query retrieved it without the language model enabled. Its ranked outputs reflected the evidence supporting clemastine but not later evidence of harm in progressive MS, which was reachable only through an on-demand literature search. Our results imply that RePurpose may be better at tracing evidence around a hypothesis than generating new hypotheses.

**Availability and Implementation:** RePurpose Pharma is freely accessible at https://www.repurposepharma.com/. For code and more information on the application, see: https://github.com/nayagamnitin-hub/RePurpose-Pharma.

## 1 INTRODUCTION

Drug repurposing is the process of finding new medical uses for existing or shelved medications that were originally studied and approved for different conditions. is important so there is not a constant chase for new drug discovery within the pharmacology field. Drug repurposing is indispensable and crucial in this field, becoming a major source of new therapies: since 2000, nearly half of the FDA drug approvals have been indications for existing products [1].

This process, in a nutshell, is done by scanning the existing databases and knowledge on pharmaceuticals to see what molecular targets they act on, whether it be agonism or antagonism, and looking at what diseases share the same molecular targets to potentially be a treatment for the disease or condition with potentially more favorable pharmacokinetics and pharmacodynamics (PK/PD). Perhaps the most notable examples of these include minoxidil and sildenafil, under the brand name Viagra. Minoxidil was originally used to treat extremely high blood pressure (hypertension) by acting as a vasodilator when researchers noted hypertrichosis, or excessive hair growth. Since then, minoxidil has frequently been prescribed in the treatment of androgenetic alopecia (or male and female pattern baldness) [2]. Respectively, sildenafil, better known by its brand name Viagra, was initially developed to treat angina pectoris and hypertension but has now been widely adopted for erectile dysfunction (ED) [3].

However, evidence is scattered across target, drug, trial, and literature databases, with separate resources and different identifiers. Connecting multiple biological targets through a multitude of different scholarly articles means crossing them all by hand. A related idea is what Swanson (1986) called “undiscovered public knowledge,” or the idea that two bodies of research are complementary to one another and bolster each other when they never cite one another. These ideas can combine to imply a conclusion that neither directly states [4]. His example in the study was fish oils with studies of Raynaud’s syndrome [5]. AI methods are becoming increasingly popular and are used across drug discovery [6], as they can help speed up the arduous process of drug repurposing and discovery.

Existing approaches utilize knowledge graphs, displaying entities including drugs, proteins, and diseases as nodes, relationships as edges, and facts stored in the databases. Some notable examples include ROBOKOP, Hetionet, ExEmPLAR, and TxGNN [7-10]. ROBOKOP uses an integrated biomedical knowledge graph built from multiple sources instead of one [7], in NCATS Translator’s effort to stop “data silos,” or the accumulation of data within one specific source rather than being spread throughout multiple sources, leading to lower accessibility of information, which NCATS aims to prevent [11]. Similarly, Hetionet integrates twenty-nine public resources to create 2.25 million relationships and score around 210,000 compound-disease pairs for probability of treatment [8]. ExEmPLAR, which stands for Extracting, Exploring, and Embedding Pathways Learning to Actionable Research, similarly uses a knowledge graph system. One unique aspect of ExEmPLAR is its ranking system, which is highly customizable by the user through a tuning system. For example, it can prioritize well-known relationships between pharmaceuticals for a specific target/disease over a range of under-researched/explored connections between the nodes [9]. This feature inspired a similar feature in RePurpose: the list carries a tunable sort control rather than one fixed order. It runs from well-established to under-explored. See Section 2.8 for more information. As deep learning has given rise to across drug discovery, there is a growing demand for open, accessible tools.

RePurpose is a web-based hypothesis-generation and literature-exploration tool that takes a written disease, target, or goal and returns candidate drugs assembled from Open Targets, ClinicalTrials.gov, UniProt, and PubMed. It combines three of the features of the above frameworks: 1) it takes free text and is thus more open to different users, 2) it assigns each target a desired direction of effect (agonist, activator, and opener count as positive; inhibitor, antagonist, blocker, and degrader count as negative) to help eliminate drugs that act in the opposite direction, and 3) it labels each candidate by its provenance (whether it is a database entry, an active trial, or a suggestion from the language model). Using the tunable ranking system, the list can also be focused from established candidates to under-investigated candidates within repurposing agents. We assess this tool with a retrospective case study of clemastine in multiple sclerosis, asking whether it recovers a repurposing hypothesis that has already been tested in a clinical trial and whether it surfaces later evidence of harm.

## 2 DESCRIPTION

### 2.1 Data Sources

### 2.2 Query Interpretation and direction of effect

When a query is first input by the user, RePurpos runs a query down one of two tracks, decided during interpretation. The first track is disease mode, in which the query names a disease (like multiple sclerosis, for example), and the biology comes from the Open Targets Platform to pull the disease-to-target associations [12]. The other track RePurpose could route the query through is the goal/targets mode, in which the query is a phenotype or goal written in plain language (eg., how to lose weight), and the model itself proposes a list of targets with a direction for each.

The model turns the free text input by the user into a JSON plan. For example, in this scenario of multiple sclerosis, RePurpose decides `mode = “disease”` and `disease_terms = [“multiple sclerosis”]`.On the other hand, if it were a *goal* query in plain language, it returns `mode = “targets”` with a list of `{symbol, direction, weight, role}`. Determining the direction the pharmaceutical needs to act in is indicated by `up` or `down`, representing whether there is more activity of the target needed to achieve the goal, such as an agonist (up), or there is less activity needed to achieve the goal, such as an inhibitor (down). There is a `weight` assigned to each based on how important that target is, from 0-1.

### 2.3 Target resolution and protein annotation

#### Disease Mode

When the query is run in disease mode, the disease inputted by the user goes directly to an Open Target Search to find the top-hit EFO ID (Experimental Factor Ontology ID). Using that EFO ID, the app pulls a ranked list of all genes (`associatedTargets`) known to be connected to that disease. For every target in that list, it pulls a package of data to be generated: `{association score, target {Ensembl ID, gene symbol, name}}`. From this list, any target that scores below 0.1 is dropped, and the top 30 targets are stored for the next step. This is the pathway through which clemastine was recommended as a repurposing agent for multiple sclerosis.

#### Goal mode

On the other hand, if the query runs in goal mode, each gene symbol the AI model generated goes to an Open Target search strictly as a target rather than a drug/disease, and an Ensembl ID is extracted for each of the gene symbols.

For the top targets that were named in either mode, UniProt turns the gene symbol into the human protein’s recommended name and function text. The source is UniProt, limited to reviewed human entries (Swiss-Prot). The report highlights the top 6 targets, and the rest are fetched on demand when a user clicks a target. This step is entirely from the direct database, so no LLM is required [13].

### 2.4 Drug retrieval and direction filtering

For each retained target (there can be up to 30), Open Targets returns every single drug that acts on it, whether it be agonistic or antagonistic on it. The tool keeps each drug’s mechanism of action and action type (agonist, inhibitor, antagonist, etc. – taken from the first listed mechanism); its max clinical stage or phase that was achieved in testing in the disease it is “known for,” or already being tested in the field of; drug type, defined by its molecular size and size; and the established safety concern, which is quantified by the adverse-event types and the count of each, black-box warnings, withdrawn flags, and warning types. Due to the fact that one pharmaceutical can hit several of these targets, the results are aggregated. In other words, the tool collects drugs that act on each of the disease’s targets, and since many drugs act on more than just one protein, it merges those answers into one entry per drug while recording two things. The targets that it reached are stored in `via_targets`, while the association score is then stored in `target_relevance`, which is used in scoring (see 2.7).

Each drug is filtered into a specific direction of action and is denoted as a number from -1 to +1. Positive numbers include agonists, partial agonists, activators, positive allosteric modulators, openers, releasing agents, and stabilizers. These scoring values range from greater than 0 to +1. Negative values represent inhibitors, antagonists, blockers, negative allosteric modulators, degraders, disrupting agents, inverse agonists, sequestering agents, and PROTACs, which range from less than 0 to -1. In the case that a direction is not recognized by the code or the action type is missing/empty, the drug is assigned a sign of 0 (neutral), meaning it neither helps nor harms the mechanism, and is effectively ignored by the directional math and scoring system.

### 2.5 Trial-derived and model-proposed candidates

Repurposing agents are grouped mainly into two groups: trial-derived candidates and entirely model-proposed candidates. Trial-derived candidates are searched from ClinicalTrials.gov for trials whose conditions match the goal. The tool extracts the drug and biological intervention and skips over non-drugs (eg., devices, exercise, etc.) and keeps the NCT IDs and the highest phase that they were tested for. A secondary measure of accuracy is enforced with a relevance check that removes any trial drugs if the trial’s listed conditions do not contain the goal term. These drugs are then marked as trial-derived by `source = “clinical_trials”` and flagged as prospective rather than established treatments and finally given scores based on the highest phase they reached in clinical trial testing.

On the other hand, model-proposed candidates are beyond the databases. The tool is given the confirmed targets with the direction (positive or negative) and the weights respectively, along with a few example database drugs for each target as an example, and thus proposes additional agents. These are explicitly labeled by the app as hypotheses through the “AI-proposed/Prospective” text on the cards for the drug. This model also has a measure of accuracy through guardrails. Each AI-proposed drug must hit one of the validated targets, or it is removed. Along with this, another directional check is enforced, removing drugs that act in the opposite of the desired effect (if the AI proposes a drug that is an agoniser of a protein that established drugs treat by antagonizing it, the drug will be dropped). Finally, names are normalized throughout clinical trials and studies to avoid duplicate cards for the same drug. One qualification of the AI-Proposed/Prospective feature is that it does tend to propose next-gen or selective agents for the same indication. An example of this phenomenon, though not inherently occurring in RePurpose’s line when testing the query “acne,” might be tretinoin versus adapalene. While tretinoin (all-trans-retinoic acid) is, and has been, established for the treatment of *acne vulgaris* (known colloquially as simply acne), it is a 1st generation retinoid that targets all three Retinoic Acid Receptor (RAR) subtypes [14], while adapalene is engineered to be selective for only RAR-beta and RAR-gamma receptors [15]. By bypassing RAR-alpha, the third subtype, it minimizes the irritation and side effects that are common in retinoids.

### 2.6 Deduplication and clinical assessment

Deduplication is ensured with both the established treatment cards and the repurposing agent cards. This is done by grouping by the active ingredient: the model groups brands/salts/formulations and even close analogs into one card. For example, all insulins are grouped into one central “insulin card.” However, similar but fundamentally different pharmaceuticals will still get their own card: while somatropin (HGH) groups together a plethora of brands such as Genotropin, Norditropin, Humatrope, Saizen, etc., long-acting forms of growth hormone, which are fundamentally different from the short-acting bioidentical growth hormone variant, are grouped under different cards (eg., Somatrogon, Lonapegsomatropin, Somapacitan, etc.). After deduplication, the drugs go into a recalibration step in which the model re-rates effectiveness and safety using its own judgment, reclassifies them as either established or repurposing agents based on the phase that they are in, removes off-goal entries, and adds well-known drugs that the database may have missed. The last feature directly uses the model’s own training knowledge, factoring them into the results, so some of the outputs may not be strictly database-derived, but the model’s prior knowledge.

Within the recalibration step, the drugs being reclassified as either established or repurposing agents is decided based on their approval stage and phase. This is a deterministic rule that runs entirely after the model and independent of it. Any drug that is not approved and is below phase 4 in testing cannot lie in the Established Treatments section and gets demoted to Repurposing Agents. This ensures investigational drugs do not wrongly get placed in the established section.

### 2.7 Scoring

Each drug is scored on a scale from 0-1 to determine the ranking and placement it is recommended to the user in. Each drug shares a `phase_norm`. `Phase_norm` = (`max_phase`)/4. `max_phase` is the phase of approval of the drug, determined on a scale from 0-4. As mentioned previously, anything under 4 is an investigational drug that is not established or approved, meaning it is going through trial phases (3/2/1), while 4 is approved, and 0 is preclinical or unknown. Drugs are calculated on an effectiveness and confidence score with different weights depending on whether the query was done in disease mode or goal mode.

#### Disease mode

There are two distinct scores generated that go into factoring for the effectiveness and confidence scores. `target_coverage = min(targets_hit / 3, 1)`. In other words, it asks how many of the disease’s relevant proteins the single drug engages, weighing drugs that touch several disease-relevant targets as more relevant than those that only touch one of the targets. After it hits 3 targets of the same disease, the score maxes out at 1.0; thus, a drug that hits only 1 or 2 can only give a partial score, while a drug that hits 3 or more of the same targets will return a value of 1.0. `target_relevance = the drug’s best Open Targets disease-target association score`. This is the drug’s strongest disease link. Of all the targets a drug acts on, it takes the highest Open Targets disease-target association score that they assign themselves. This value ranges from 0-1 and measures how strongly a protein is tied to the disease. As for the two main pillars: `effectiveness = 0.6·phase_norm + 0.25·target_coverage + 0.15·target_relevance and confidence = 0.7·effectiveness + 0.3·target_relevance`. These are the two scores the drugs are genuinely scored on, along with the distinct weighting of each variable for the scoring system in disease mode. The weighting, as well as the variables used, differs in the goal/targets mode.

#### Goal mode

In this section, the two distinct variables/scores that factor into the effectiveness and confidence scores are `goal_fit` and `establishment`. `goal_fit = clamp(best_help + 0.1·(n_help−1) − 0.5·best_harm, 0, 1)`.Ranging from 0 to 1, this value is indicative of how well the drug pushes your targets the desired way. It is built from `best_help`, the largest weight among the targets it pushes in the right direction; `best_harm`, the largest weight it pushes in the wrong direction; and `n_help`, how many targets it helps. `Establishment`, on the other hand, is a binary property: `establishment = 1.0`if the drug is already established for the goal; otherwise, it is 0.45. As for the two main pillars of weighting and scoring: `effectiveness = 0.5·goal_fit + 0.2·phase_norm + 0.3·establishment` and `confidence = 0.6·effectiveness + 0.4·goal_fit`.

In both models, they utilize a `safety` score. The values for each type are shown below, respectively.

Most importantly, every drug’s score is computed twice, with the second pass winning. The first stage of scoring is through the formulas mentioned above, stored as `_score` in pipeline.py. This uses the database facts to assign the first-draft `effectiveness, safety`, and `confidence` scores.

In the second stage, or the assessment pass (stored as `assess_and_augment` in assess.py), the model re-rates each drug’s effectiveness and safety from its own clinical knowledge and overwrites the Stage-1 numbers (if a live model is connected). The assessment stage’s formulas are listed below:

~~~
def _apply(c, d): # d = the model’s rating for this drug
  eff = _clamp(d.get(“effectiveness”), c.effectiveness_score)
  safe = _clamp(d.get(“safety”), c.safety_score)
  c.effectiveness_score = round(eff, 3) # ← REPLACES the formula’s value
  c.safety_score = round(safe, 3) # ← REPLACES the formula’s value
  c.confidence = round(0.7*eff + 0.3*(c.goal_fit or eff), 3) # ← recomputed from the model’s eff
~~~

Thus, the final confidence score weighting is 70% and 30%. On a normal run, with the model on, the formula numbers exist for a split second but are then erased quickly by this line of code: `c.effectiveness_score = …` Therefore, the heuristic formulas only serve as a fallback when the model is off or when the call fails, and scores reported are the model-recalibrated values. One thing to note is that not every single drug necessarily goes through this assessment pass second stage. The assessment pass caps how many it scores at only around the top 45, and a long tail keeps their formula scores.

### 2.8 Tunable ranking

This feature was inspired largely by ExEmPLAR (Extracting, Exploring, and Embedding Pathways Leading to Actionable Research), a biomedical research database consisting of knowledge graphs. ExEmPLAR directly features a “highly tunable ranking system” that allows the user to “prioritize strongly supported relationships with numerous co-mentions or under-explored relationships with few co-mentions” [9]. Co-mentions, in this case, mean the scientific literature in studies backing them up. RePurpose draws directly from this – the repurposing agents have a slider that the user can control to prioritize either well-established/high evidence drugs or early/under-explored drugs. The user can put the slider anywhere in the range. The ranking system will only reorganize, not redo the whole query again. Thus, the user must click the “Sort” button before it is resorted to their choosing. The formula for the ranking system is based on how established a drug is and its novelty, which is factored into a score. The source weight has three fixed options, which it could possibly be: `source_weight`: model-proposed (`ai`) = 1.0, trial-only (`clinical_trials`) = 0.6, database = 0.0. Novelty is determined based on the source weight: `novelty = 0.5·(1 − phase/4) + 0.5·source_weight`. The earlier stage the drug is in its development and the more model-generated it is, the more “novel” the drug is considered. Finally, they are given display scores: `display_score = α·novelty + (1 − α)·confidence, sorted descending`. When α = 0, the model has the highest confidence. The drug is well-established, and there is strong evidence for it as a repurposing agent already. On the other hand, if α = 1, it indicates pure novelty. The drug is early and underexplored.

### 2.9 Evidence retrieval and conversational assistant

For each drug card shown, whether it be a repurposing agent or an established treatment, there are two buttons shown to the user: view studies and ask AI. The View Studies button retrieves the PubMed studies for that specific drug and the topic that is searched via NCBI E-utilities, and the article titles are fetched and displayed to the user. Every link is accessible and shown as a hyperlink. The other button, the Ask AI feature, is specifically on drug cards to act as a focused Q&A. The system will retrieve PubMed snippets for the drug, then answers the user’s question as well as cites the PMIDs for anything drawn from the studies and ensures to label mechanistic inferences as “Mechanistic reasoning.” Additionally, there are symbols on each card that denote special characteristics about each one (e.g., black-box warning, most effective, etc.) When clicked, a short description tailored to the specific label will explain what the label means. Finally, protein target chips can also be clicked to give a short description of what the target protein does through a UniProt lookup. These protein target chips appear on the screen after a query is made as well as on specific drug cards.

Another separate layer of an LLM is available through a conversational assistant called the “Personal AI Pharmacologist.” This is entirely separate from the structured results and acts as a chat agent to answer free-form questions. It runs through an n8n workflow running Claude Sonnet 5, and thus the browser calls on it during its chat. It can fetch and read any paper it is given, whether it is given to it in the form of a PMID, DOI, or PMCID link. This is the key property for the case study, as this answers the model’s parametric knowledge and is not bound by the disease versus goal/target modes in the query section. The most interesting aspect of this conversationalist assistant is the fact that it can surface off-target repurposing candidates (like clemastine) that the structured pipeline could potentially miss in disease mode.

### 2.10 Implementation

The active model that RePurpose Pharma uses is Claude Sonnet 5 (`anthropic/claude-sonnet-5`), which is accessed through an OpenRouter API Key. This is selected by configuration. Each result from a query is cached as a lower-cased key in a bounded store so the same query returns the same report. However, a refresh path does exist to force a fresh recompute and ignore the saved copy. Since this pipeline is so stochastic, this cache system ensures reproducibility within the system. In the scenario that the AI cannot be reached, there is a backup plan in place. The tool will still run, but without all of the AI parts. The non-LLM mode still passes query interpretation via a hand-written keyword map or a literal disease search, and can access databases and clinical-trial candidates through Open Targets and ClinicalTrials.gov, and can access structural formula scores. However, the AI mode is still needed for the model to propose candidates (AI-Proposed/Prospective drugs) and for the canonicalization, recalibration, and summary processes. Without being able to access the AI (in the case that there is no key, the model is down, or it is simply out of credits), drugs cannot be grouped based on brands or salt variants and cannot be recalibrated through the AI re-scoring system. This also inherently means the conversational AI will not work.

## 3 Case Study

### 3.1 Background

Immune-mediated inflammatory damage to myelin leads to a chronic central nervous system disease called multiple sclerosis (MS). Despite the fact that no disease-modifying therapies have yet been approved specifically to repair the myelin in MS, the available therapies show a potential anti-inflammatory effect without repairing the already damaged myelin. This implies that remyelination persists as an unmet therapeutic goal [16].

Clemastine fumarate, a first-generation H1 antihistamine drug available by prescription and over the counter (OTC) [17], has been used as an allergy suppressant since 1977 [1]. It was identified as the top-performing drug for MS during a high-throughput screening of bioactive agents that stimulate the differentiation and myelinating potential of oligodendrocyte progenitor cells [18]. This was later ascribed to blocking the M1 muscarinic acetylcholine receptor (CHRM1). This action was potentially off-target and not a consequence of its primary action at the H1 histamine receptor [19].

Clemastine was later tested in the ReBUILD trial, a monocentric, double-blind, randomized, placebo-controlled crossover phase 2 trial (50 patients) for relapsing MS and chronic demyelinating optic neuropathy [20]. A 150-day course of treatment was shown to decrease the visual evoked potential latency delay by 1.7 milliseconds per eye, which could represent remyelination on the visual pathway, though no significant difference in clinical outcome was detected. Clemastine thus constitutes a genuine case of drug repurposing: an approved medication for a different indication, having both mechanistic rationale and clinical evidence for use in a new indication. However, its safety in progressive MS was challenged as described in Section 3.4.

### 3.2 Query protocol

To assess whether RePurpose supports the clemastine hypothesis, we ran the tool three separate ways. Firstly, in disease mode, RePurpose was asked “multiple sclerosis” and “demyelinating disease”. Next, in goal mode, RePurpose was asked “promote remyelination in multiple sclerosis” and “remyelination / oligodendrocyte precursor differentiation,” for which it provided an indicated target and direction of effect for each. Lastly, we asked the RePurpose conversational assistant two questions about candidate repurposes for remyelination (each in a separate chat window). The exact text of all of the prompts and questions is in Supplementary File S1.

All experiments were performed on 2026-10-13 with the following parameters: Claude Sonnet 5 (anthropic/claude-sonnet-5) accessed via OpenRouter, code version for RePurpose= 81f67e4 (2026-10-10). Each structured query was run once with the result cache disabled to recompute the entire pipeline (runtimes between 250 and 430 seconds). The candidate-proposal step was performed at a sampling temperature of 0.2, which implies that the set of model-proposed candidates can differ between runs. Re-purposed candidates were ranked at the default slider position (α = 0).

During each execution, data on whether clemastine was found, the list the clemastine was found in, the position in the list and the length of the list, the label of the list, the source provided; the mechanism and targets provided, the proficiency, safety, and confidence scores, the trial IDs that were linked to, and whether the resolved targets included the H1 histamine receptor (HRH1) or the M1 muscarinic receptor (CHRM1). Full reports are provided in Supplementary File S1.

### 3.3 Recovery of clemastine

Table 3 lists whether clemastine was found successfully in RePurpose for each query, which varied depending on whether the disease or disorder mode was used. In disease mode, clemastine was not returned as a result for either “multiple sclerosis” or “demyelinating disease”, as the targets returned from Open Targets in both disease conditions were almost exclusively immune- and inflammatory-related proteins, including NR3C1, MS4A1 (CD20), sphingosine-1-phosphate receptors, integrins, interferon receptors, and potassium channels. The muscarinic receptors CHRM1 and CHRM3 were not among the top-30 highest-scoring targets, which were all above the 0.1 association-score cutoff, and so not accessible via the target route for this mode, despite clemastine’s mechanism of remyelination being a consequence of its antagonism of CHRM1 [19]. In addition, the clemastine therapeutic was not retrieved when searching for the clinical trials, as this search would only return a handful of trials per condition and only identify the top-scoring results for each condition term. For the broad disease condition, the top results were almost all large trials of disease-modifying therapies, which excluded small, remyelination-specific trials of clemastine.

**Table 1.**
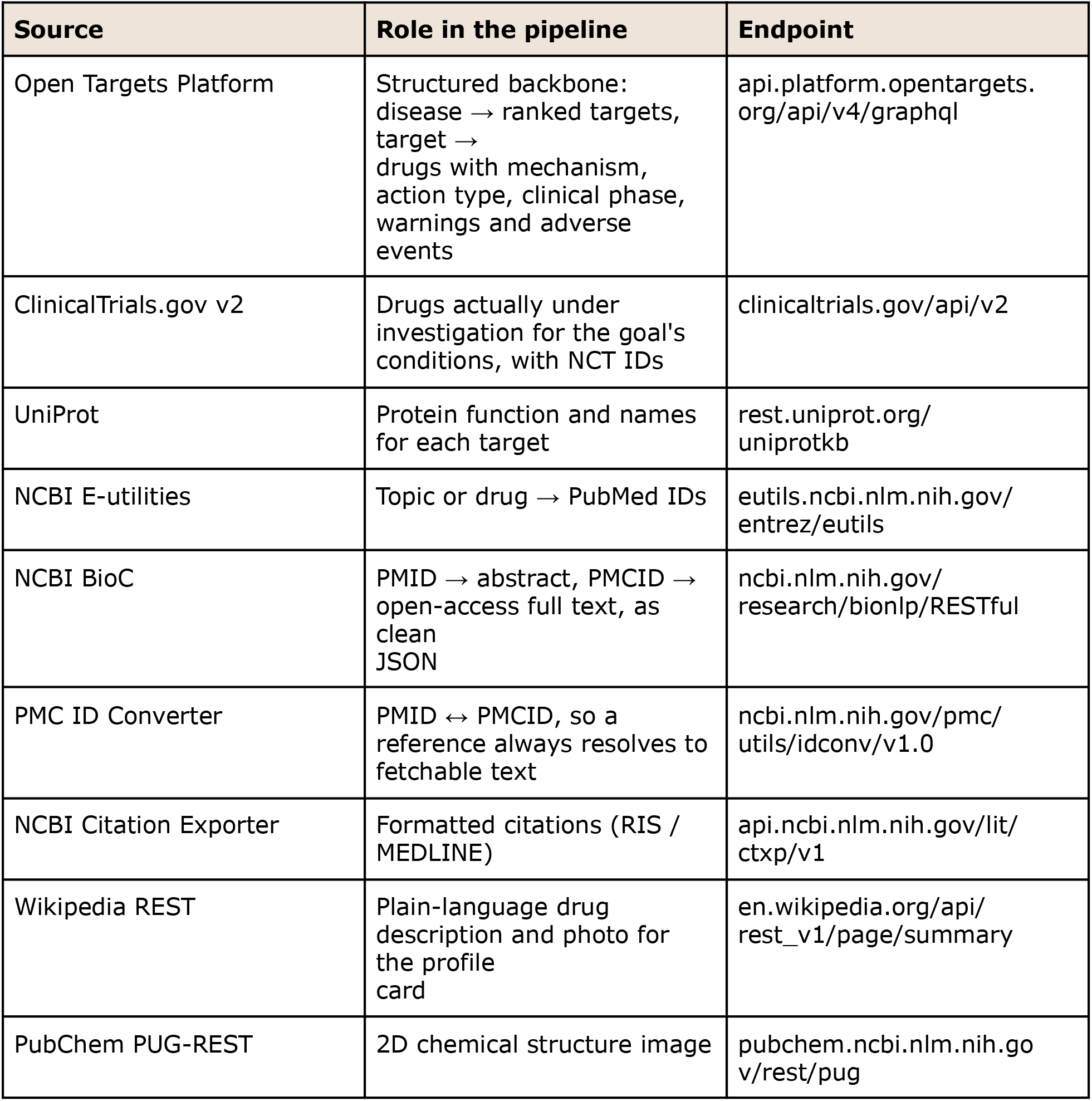

**Table 2.**
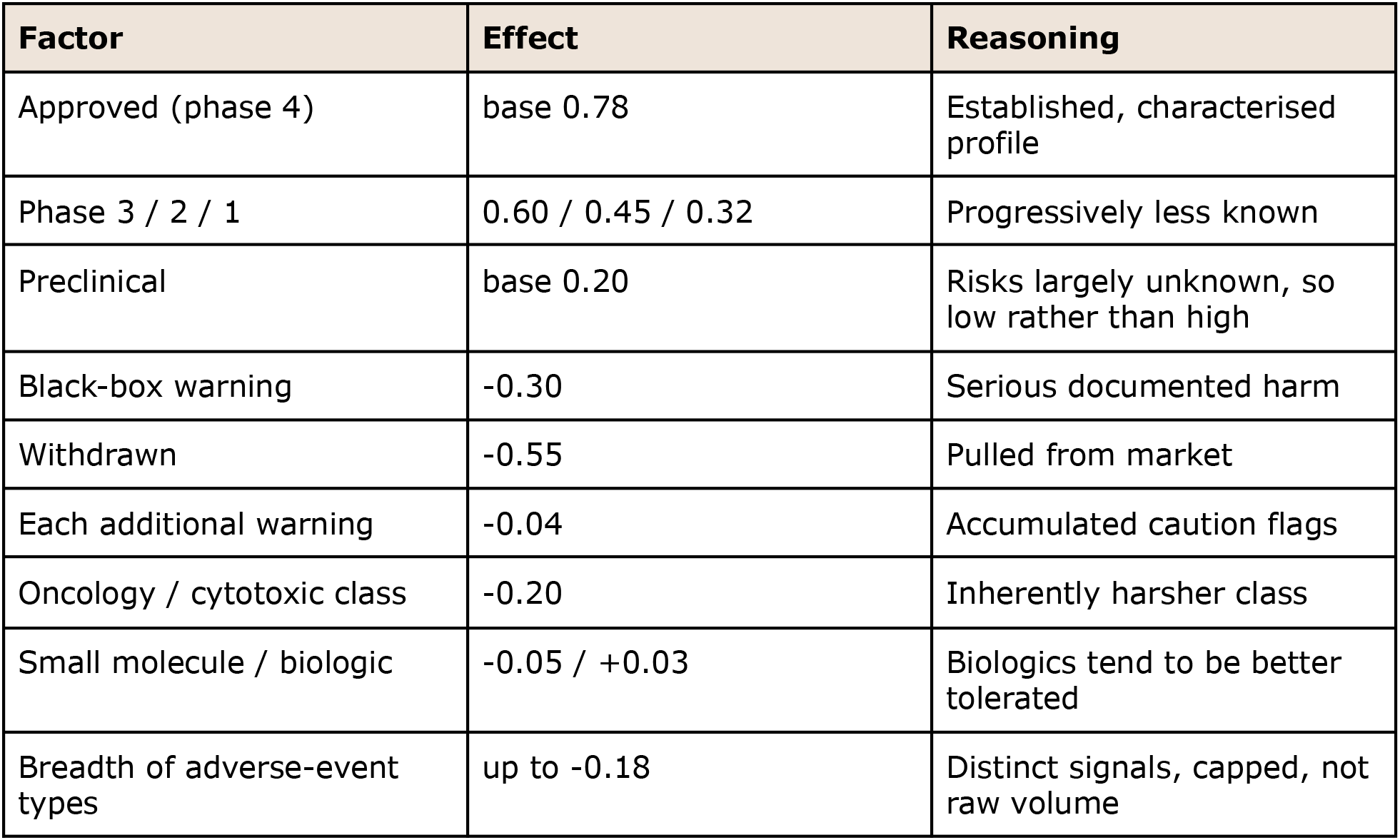

**Table 3.**
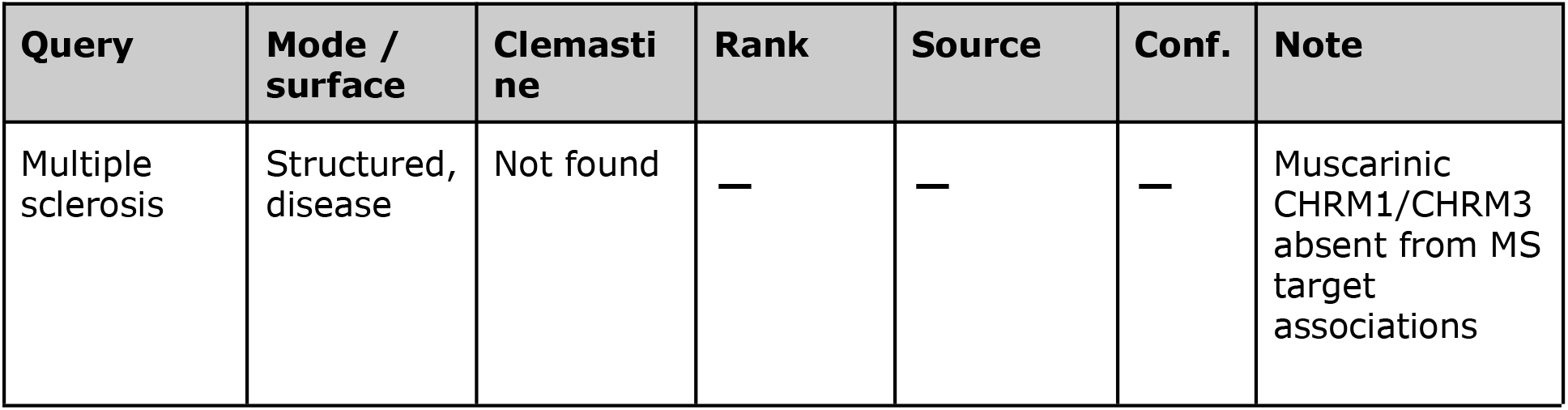
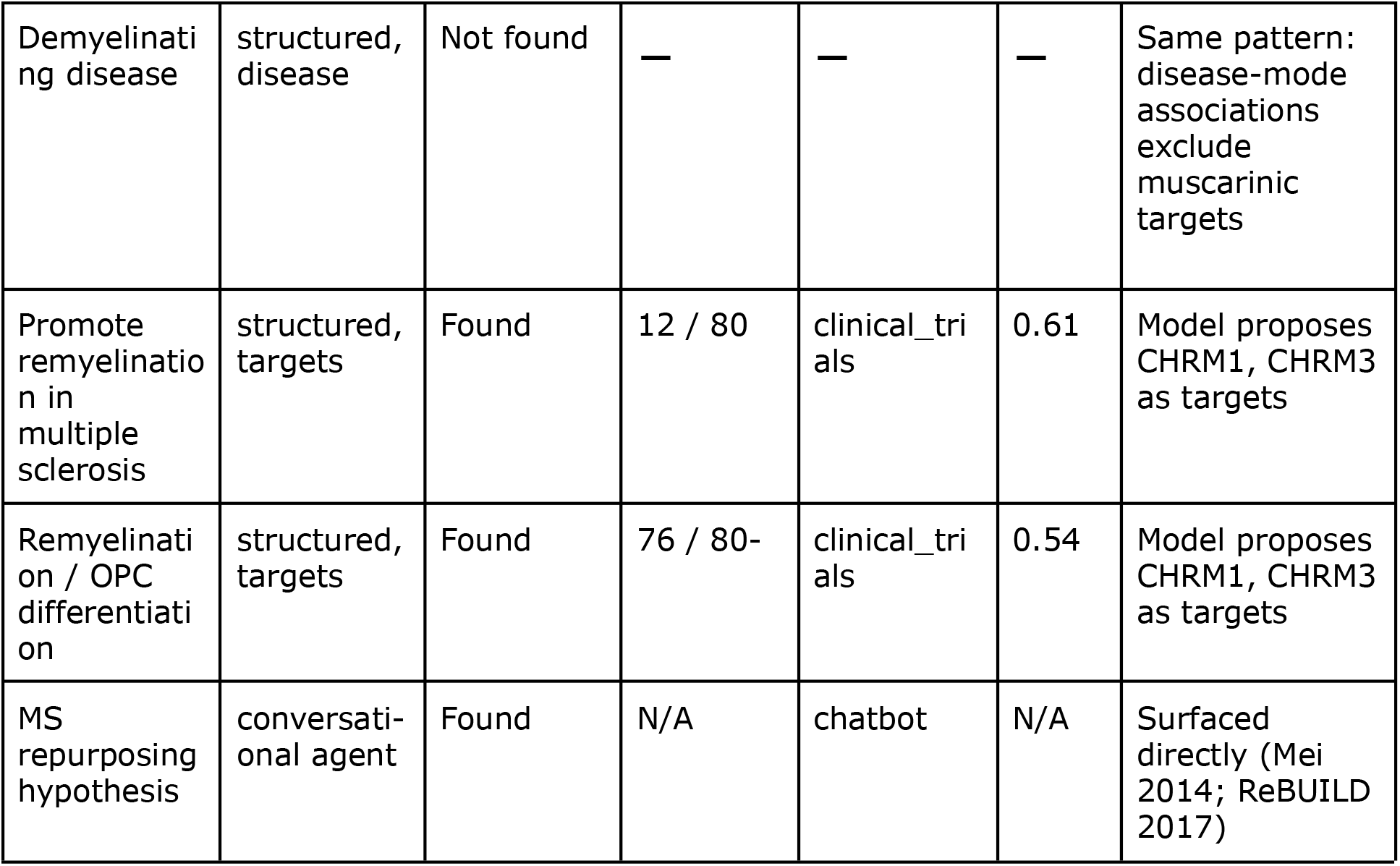

| Query | Mode / surface | Clemastine | Rank | Source | Conf. | Note |
| --- | --- | --- | --- | --- | --- | --- |
| Multiple sclerosis | Structured, disease | Not found | — | — | — | Muscarinic CHRM1/CHRM3 absent from MS target associations |
| Demyelinating disease | structured, disease | Not found | — | — | — | Same pattern: disease-mode associations exclude muscarinic targets |
| Promote remyelination in multiple sclerosis | structured, targets | Found | 12 / 80 | clinical_trials | 0.61 | Model proposes CHRM1, CHRM3 as targets |
| Remyelination / OPC differentiation | structured, targets | Found | 76 / 80- | clinical_trials | 0.54 | Model proposes CHRM1, CHRM3 as targets |
| MS repurposing hypothesis | conversational agent | Found | N/A | chatbot | N/A | Surfaced directly (Mei 2014; ReBUILD 2017) |

Conversely, clemastine was proposed for both of these queries in goal mode. For each, the model proposed CHRM1 and CHRM3 as targets of inhibition (entry in results from ClinicalTrials.gov where clemastine appeared as a result from source NCT05338450, or from the metformin-clemastine combination trial NCT05131828 [21], instead of as a model-proposed candidate). The model-proposed candidates for the two queries differed: clemastine was ranked 12/80 of repurposing candidates (confidence score 0.61) for “promote remyelination in multiple sclerosis”, but 76/80 (confidence 0.54) for “remyelination / oligodendrocyte precursor differentiation”. The model-proposed candidates for these two queries included other candidates for muscarinic signaling or remyelination, such as telenzepine, PIPE-307, and clobetasol, though clemastine did not.

In addition, the conversational assistant directly noted clemastine as a repurposing hypothesis for multiple sclerosis (original screen [18] and ReBUILD trial [20]). Since both this response and the selection of the muscarinic targets in goal mode are based on the LM training data, these results indicate the tool’s ability to retrieve the hypothesis instead of discovering it (discussed further in Sec. 4.2).

### 3.4 Contrary evidence

Three studies using ReBUILD have added more evidence to the clemastine hypothesis. The CCMR-Two trial was a randomized, placebo-controlled phase 2 trial with 67 patients with relapsing-remitting MS on a stable disease-modifying agent and assessed the effect of combination clemastine and metformin; the primary endpoints were compatible with remyelination on visual evoked potentials [22]. However, the combination of metformin with clemastine and the presentation of this study as a conference abstract preclude it from isolating the specific effect of clemastine. Meanwhile, in the TRAP-MS platform trial in patients with progressive MS, patients randomized to a clemastine arm experienced a faster rate of disability accumulation, and the arm was closed when 3 of the 9 patients accumulated enough disability to meet safety stopping criteria [17,23]. The investigators linked this to clemastine enhancing pyroptosis, an inflammatory form of cell death, in macrophages and oligodendrocytes exposed to extracellular ATP [17]. In a rabbit model of chronic demyelination, which was chosen over rodents because its oligodendrocyte progenitor cell densities are closer to MS lesions, prolonged administration of clemastine increased generation of oligodendrocytes but reduced the progenitor pool, and delayed treatment in the first month produced progenitors with senescence markers [24]. Taken together, these studies imply that clemastine may be effective at certain points in the disease process and in certain durations of treatment.

The ranked results and conversational assistant of RePurpose largely did not include this contrary evidence. In the goal-mode runs in which clemastine was presented, it was mapped to registered remyelination trials, and its ranked report failed to include the TRAP-MS publication or the rabbit experiment. Its safety score of 0.70 might have been due to its status as an approved drug, not the AD stage diseases (progressive MS) in which it failed, and its conversational assistant read only from the initial screen and ReBUILD [18,20]. Its only exception was in the on-demand literature search: the PubMed studies returned for clemastine via View studies had the TRAP-MS disability paper at the top of the results set, though not the rabbit-model paper, and only if the user chose to look at that view. This likely reflects the nature of the pipeline: the trial source records specify only that a trial exists and was run in a phase, not how it fared or ended, and the safety score does not include indication or stage of disease. Consequently, rankings and scores presented the evidence for clemastine and relegated that against clemastine to an information retrieval operation the user has to perform, a limitation we discuss in Section 4.2.

### 3.5 Effect of tunable ranking

To demonstrate the influence of the tunable ranking on a known repurposing candidate, the order of the repurposing candidates returned for the goal-mode queries was recalculated at five slider values (α = 0, 0.25, 0.5, 0.75, 1.0) using the scores from the runs reported in Table 3. Clemastine entered from the trial source via a source weight of 0.6 and clinical phase of 3 with a novelty score of 0.425. Its rank at each slider position is shown in Table 4. For the query “promote remyelination in multiple sclerosis,” clemastine was 15th out of 80 rankings (default order) at α = 0, elevated to 9th, fell back to 12th, and then moved lower to 21st. For the query “remyelination / oligodendrocyte precursor differentiation,” clemastine ranked 76th, out of 80. The movement was non-monotonic: at lower novelty weightings, a distribution of approved drugs ranked higher-confidence than clemastine; above clemastine dropped to zero novelty, temporarily raising clemastine; then at higher weightings, the earliest stage and model-proposed candidates, with more novelty than a phase-3 trial drug, overtook clemastine. This demonstrates the control system’s ability to raise an authentic, trial-stage repurposing candidate above more established options at moderate novelty weightings. Its shortcoming is that a drug, like clemastine, that has already reached phase 3 will be knocked down at higher novelty weightings, so the control elevates clemastine only over a moderate range of novelty weighting, not over the extreme “under-explored” range.

### 3.6 Repeatability and contribution of the language model

To evaluate repeatability, we ran each structured query 5 times with the result cache disabled. In disease mode, clemastine was not present in any run of either query. In goal mode, clemastine was present in 5/5 runs of each query, but the candidate-suggestion rank and origin class differed from run to run; for “promote remyelination in multiple sclerosis”, candidate-proposal rank ranged from 15th to 55th of 80, and candidate-origin class ranged from clinical-trials candidate (clinical phase 3) (3 runs) to approved database entry (phase 4) (2 runs); for “remyelination / oligodendrocyte precursor differentiation”, candidate-proposal rank ranged from 6th to 33rd of 80, and candidate-origin class was clinical-trials candidate (clinical phase 3) (4 runs) to database entry (phase 4) (1 run). The model-proposed candidates also varied: telenzepine and PIPE-307 appeared in 4 and 1 of 10 goal-mode runs, respectively. This variation is not unexpected, because the candidate-proposal step samples from a distribution with temperature 0.2, and the assessment pass that rescored, reclassified, and reordered candidates uses the model.

We also repeated the structured queries with the language model turned off. In the absence of a model, goal-mode queries reverted to a literal disease interpretation of the query string, which didn’t match any disease and resulted in no candidates, while disease-mode queries returned only database and trial candidates and the structural scores with which they entered the candidate pool, and did not list clemastine for any query. Hence, clemastine was recovered by the structured pipeline only via the path through the language model when the latter interpreted the query by mapping the mechanism to muscarinic targets, thus routing the query into goal mode, and by providing the condition terms in the trial search through which clemastine entered. Because the conversational assistant is implemented by the same model, both routes through which RePurpose recovered clemastine depend on that model, whose training data likely include the published clemastine studies. We talk more about this limitation in section 4.2.

## 4 Discussion

### 4.1 Interpretation of the case study

The case study posed two questions: does RePurpose discover a prior clinical-trial-validated repurposing hypothesis? And does it reveal additional evidence of harm? (Section 1). The answer to the first question is partly yes. RePurpose retrieved clemastine in all goal-mode runs, but not a single disease mode run (Section 3.3, 3.6). The answer to the second is almost entirely no. The ranked list of results and their scores are consistent with the supporting evidence, and the evidence of harm was only accessible through an on-demand literature search (Section 3.4).

It is not a random consequence of the failure of a disease-mode search; it is a property of what the search covers. Open Targets doesn’t simply hold up one treatment for MS; it ranks the top-ranked targets for MS by the strength of the evidence connecting them to MS [12]; it is thus no surprise that the top-ranked mechanisms are all to do with how the immune system goes wrong in MS. Clemastine’s proposed mechanism is different: to repair myelin, it works through the oligodendrocyte precursor cells, by blocking the M1 muscarinic receptor [18,19]. The disease-to-target search describes what is known about how a disease goes wrong, not how it can be repaired, and so a candidate working through anything that is not part of the disease’s disease-to-target profile would be unreachable on the disease side. The trial route failed for another reason: it only looked up to the top few trials for a given condition and for MS, being so well studied, those turned out to be established therapies. Both failures matter, because candidates that act through a mechanism unrelated to a disease’s known biology are precisely the cases repurposing tools are meant to find.

The answers were different when the question was posed as a mechanism: in goal mode, the language model committed the same query, muscarinic targets, and the source language search term (Section 3.3), which was precise enough to bring back the remyelination trials, and entry of clemastine into the candidate list with its trial identifiers and a source label. This should be read as recovery rather than discovery. With the model disabled, no structured query returned clemastine (Section 3.6), and the conversational assistant, which runs on the same model, named clemastine while citing the original screen and the ReBUILD trial [18,20]. Very probably, the model was trained on these articles, and its association of remyelination with muscarinic is memorized, not derived [25]. Same old contamination, in other words. But what the demonstration does show is that the pipeline can convert a mechanism-level question into a ranked candidate with traces of its sources that can lead back to registry entries and then back to articles, just as the instrument was intended to.

Clemastine’s position in the list was far less predictable than its presence, as it was inconsistent across runs, whose single query locations (Table 3) ranged from 12th to 76th of 80, with sources switching between trial and database (Section 3.6). While the tunable rankings pushed the clemastine higher at more moderate weightings of novelty, at the extreme, the drug was ranked lower, because the novelty measure treats a trial-stage drug as less novel than early-stage and model-proposed candidates. A candidate’s rank in a single run is therefore a weak signal of its plausibility, and the list is better read as a set of hypotheses to examine than as an ordering to trust.

The most consequential result concerns the evidence against clemastine. The record now shows its effect is context-dependent: a remyelination signal on a biomarker in relapsing MS [20], a signal from combination treatment in CCMR-Two [22], accelerated disability in progressive MS [17,23], and progenitor exhaustion with prolonged treatment in a rabbit model [24]. RePurpose’s scored outputs reflected only the first part of this record. For example, because the trial source only records that a trial exists and its phase, and not its outcome, it cannot distinguish a trial that reported a benefit from one that stopped for harm; its safety score is not sensitive to indication or disease stage. The harm was not suppressed: the TRAP-MS paper was the top result returned by View studies (Section 3.4). But the ranked list and scores would have concealed it from a user. In Swanson’s terms [4], RePurpose retrieved the literature supporting the hypothesis while ignoring those that contradict it. With a hypothesis-generation application, retrieving disconfirming evidence is important as well as retrieving supporting evidence, a point we revisit in Sections 4.2 and 4.3.

### 4.2 Limitations

The tool is evaluated using only one retrospective case. Choosing Clemastine in MS as the case study was primarily due to its well-researched history of repurposing, which provides a test as to whether the tool can rediscover a known hypothesis. One pair of drugs and indications cannot provide an estimate of how often RePurpose can rediscover hypotheses or how often it ranks failed ones highly in general. These pairs of approved and failed drug-indication pairs already exist to this end [26], and RePurpose has not been compared with existing drug-repurposing tools [8-10] using any shared task. Our results are also only reflective of the state of the underlying databases at the time of the runs, and may change as Open Targets and ClinicalTrials.gov are updated.

Second, the experiment is not necessarily independent of the training data of the language model. Clemastine’s remyelinating effect, its muscarinic mechanism, and the ReBUILD trial were all published years before current language models were trained [18-20], and every route by which RePurpose recovered clemastine passed through the model (Section 3.6). The recovery may therefore reflect recall of published findings rather than an ability to generate new hypotheses [25]. The question format compounds this. The goal-mode queries that recovered clemastine named remyelination and oligodendrocyte precursor differentiation, which is already the mechanism behind the hypothesis, and they were designed with knowledge of the expected answer. A fair test would require hypotheses established after the model’s training data were collected, queried by someone blind to the expected answer.

Third, the retrieval design restricts the set of candidates that can be retrieved. The pipeline links drugs to a disease in a single step, through targets already associated with that disease, so it cannot reach candidates that act through a mechanism outside that association profile, as clemastine does through CHRM1 [19]. Multi-hop graph methods are designed for these indirect connections [8,10]. Open Targets association scores also reflect where evidence has accumulated, which in this case favored immune targets over repair biology [12]. Separately, the trial search retrieves only a few top-ranked trials per condition, so small or mechanism-specific trials are excluded for heavily studied diseases.

Fourth, the scores are heuristics, and the ranking is not stable. The weighting in the scoring formulas was chosen by hand, without outcome validation. With the model linked in, the assessment pass replaces the formula scores for approximately the first 45 candidates with the model’s own ratings, and keeps the formula scores for all following candidates, so a single ranked list contains two separate scoring regimes that are not on the same scale. As the proposal and assessment steps sample over the model, clemastine’s rank, and even the label of its source, changed between five runs that started from the same seed (Section 3.6). Caching stabilizes a specific saved report, but not the pipeline that produced it, and five runs are too few to reliably estimate that variation. Additionally, the measure of novelty used to determine tunable ranking disfavors approved and late-stage drugs, despite approved drugs representing the very class of compounds that are re-used in any repurposing effort. This explains clemastine’s drop-off at the highest novelty weightings (Section 3.5).

Finally, RePurpose does not consider evidence against a candidate. Its trial source indicates only that a trial is documented and the phase maximum. The safety score is independent of the outcome of each trial, since it does not consider trial failures or early stops (for harm), only the existence of a trial. Nor does the safety formula or the model’s re-rating take indication or disease stage into account, so clemastine received a safety score of 0.70 despite evidence that it accelerated disability in progressive MS [17,23]. This negative evidence was available via View studies (Section 3.4), but only to a user who looked for it. The conversational assistant shares this limitation, and its answers were not evaluated systematically, since we asked it only two questions.

### 4.3 Responsible use

RePurpose is meant for researchers, students, and members of the public interested in pharmacology and considering how and whether existing drugs could be repurposed. It is a hypothesis-generation and educational resource, not medical advice, and should not be used to inform treatment or self-medication. Since it is available to the public and the assistant speaks in everyday language about goals (including weight loss and regrowth of hair), there is the potential for a user to interpret a ranked candidate as a recommendation, and the case of clemastine illustrates why that is problematic. A drug that has been for sale over the counter for decades, which in one trial showed a remyelination signal [20] and in another patient population was linked to faster disease progression [17,23] and was hard for the RePurpose ranked output to pick up harm on (Section 3.4). Availability does not mean a drug is safe for a new indication [17].

Several protections are built in. There is a disclaimer on the banner: research only, not medical advice. Candidates from the trial and model are shown as “prospectives,” and model-proposed as well as trial-derived candidates as “prospective,” so no one should mistake hypotheses for treatment. A rule is used to avoid a drug that has not gone through phase 4 appearing among identified treatments (2.6). The drug cards showed the registry status of linked trials and linked claims to studies in PubMed so one could verify the evidence directly. The conversational assistant was limited to topics in pharmacology and research, and before it would recommend that a user add, combine, or swap a drug, it was instructed to state any clinically significant interactions with any drugs mentioned by the user, and to say when a question needs a clinician. These safeguards reduce but do not eliminate misuse; the interaction rule is an instruction, not a well-founded database, and no label or disallowed banner can prevent anyone from acting on a candidate.

Hence, we advise that individuals use each output as a prompt for reading rather than a definitive answer. Individuals should verify the status and outcomes of any related trial, View studies for counterevidence as well as for a candidate, and seek advice from a family member’s physician or pharmacologist prior to taking, ending, or changing any type of medication. They also need to know that what is typed into the conversational assistant (such as medicines that they might tell the conversational assistant that they use) is handled by third-party services (OpenRouter and an n8n workflow) and that the assistant associates the memory with a session identifier that resets when the user restarts the chat.

### 4.4 Future work

The next, most critical step is an objective evaluation. An approved and failed drug-indication-pair benchmark (e.g., repoDB [26]) would quantify the proportion of RePurpose confirmed repurposings, and of failed cases ranked highly, rather than a single instance. To distinguish RePurpose’s reasoning from language-model recall [25], such a benchmark would consist of repurposings approved or published after the collections of language-model training data, and the queries written by authors who do not know the answers. Such a benchmark would also make it easy to compare with knowledge-graph and graph-learning tools [8-10], conduct more repeat runs for measuring run-to-run variation, and test whether a lower sampling temperature, or combining multiple runs, yields more stable rankings.

Second, evidence against a candidate should count against it. Drug cards already state the registry status of the trials they are in; the next step is to use the status when scoring so that a failed or terminated-for-safety trial, for example, reduces the candidate’s confidence rather than raises it. Registry status alone would not have tipped us off on clemastine, since the TRAP-MS trial only terminated its clemastine arm, not the entire trial [17,23]. The pipeline should also link to published safety and efficacy results for each candidate and indication, and list them on the card rather than expect a user to find them on demand. The safety score should also become indication-specific and, when evidence warrants, disease stage-specific.

Third, retrieval might identify candidates that the current design overlooks. Paginating the trial search, or ranking trials by relevance to the mechanism of the query instead of stopping at the top few per condition, would retrieve small, mechanism-focused trials for heavily studied diseases. The case study indicates that combining the two modes of query is feasible: for the disease query, the pipeline could be asked to also suggest candidates at the mechanism level, e.g., targets involved in tissue regeneration, and mark any candidates found this way as model-derived. This would have let clemastine be retrieved from the query “multiple sclerosis” while preserving the provenance. Multi-hop connections similar to those used by graph-based methods [8,10] present another possibility that does not depend on the model.

Last, scoring and ranking. The formula weights should be scaled relative to the benchmark, and the formula and model scores should be recalibrated or applied to each candidate to enable ranking on the same scale. The novelty score should consider the drug’s phase for the queried indication, not overall development, so that approved drugs being tested for a new use are not treated as already established. The assistant should be evaluated in a standardized way using a fixed set of questions with human raters knowledgeable in pharmacology.

## 5 Conclusions

RePurpose searches for repurposing candidates that answer plain-language research questions by leveraging public sources of target, drug, trial, and literature data, and cites each candidate to its source for ease of investigation. RePurpose learned to identify clemastine as a candidate for remyelination in multiple sclerosis when the question was presented as a mechanism, but not as a disease, due to the influence of the language model, which likely learned from the literature on which the model was trained, including the clemastine literature. The tool’s scored outputs tracked with evidence supporting clemastine but missed the evidence of potential harm later identified in progressive MS, which could only be retrieved by searching the literature on-demand. The results suggest that a tool of this sort is useful for hypothesis organization and validation, rather than hypothesis discovery, and that evidence weighing of the candidate is just as important as evidence retrieval for a candidate. Next steps are to evaluate RePurpose on a benchmark of approved and failed repurposings, in which the failed examples were published after the model’s training data were prepared.

## Data and code availability

RePurpose is freely available at https://www.repurposepharma.com. Its source code is available at https://github.com/nayagamnitin-hub/RePurpose-Pharma, and the case-study runs used commit 81f67e4 (10 October 2026). All data sources queried by the tool are public (Table 1). The query strings, conversational-assistant prompts, and reports from the case-study runs are provided in Supplementary File S1.

## Author contributions

N.N. conceived and developed RePurpose, designed and performed the case study, and wrote the manuscript. A.A. supervised the project, provided guidance on the scientific approach and the manuscript, and reviewed and edited the manuscript. Both authors approved the final version.

## Competing interests

N.N. developed and maintains RePurpose and the repurposepharma.com website. A.A. is employed by Sirpi Products and Services Private Limited, where this work was carried out as an internship project. The authors declare no other competing interests.

## Funding

This work received no external funding. Model API costs (Claude Sonnet 5, accessed through OpenRouter) were paid personally by N.N., and the internship during which RePurpose was developed was unpaid.

## Acknowledgments

We thank Brijesh M. and Nithin for building the n8n workflow behind the conversational assistant, and Anand Lakshmanan (Sirpi) for facilitating the collaboration between the authors. We also thank Sirpi Products and Services Private Limited for hosting the internship during which RePurpose was developed.

## Use of AI tools

RePurpose employs a large language model (Claude Sonnet 5, Anthropic, via OpenRouter) for query interpretation, candidate suggestion, candidate evaluation, and its conversational assistant (Section 2). The RePurpose code was created primarily with Claude Code (Anthropic) (used within VSC), a commercial AI coding assistant; N.N. designed the tool, guided its creation, and performed its behavior testing. Claude Code was also used to run the case-study queries, produce the analyses reported in sections 3.3 to 3.6, and create Supplementary File S1. The Figure 1 layout and tabular content were generated with Claude Code from the tool outputs and redrawn by the authors in Google Drawings and Google Docs. The authors used Claude (Anthropic) to suggest a paper structure, identify and verify sources, write example paragraphs, and review drafts for accuracy and clarity during preparation of this work. The authors have rewritten and checked all text. The authors take full responsibility for the contents. Supplementary File S2 was assembled with AI help on the tool documentation and verified by the authors.

**Figure. 1.**
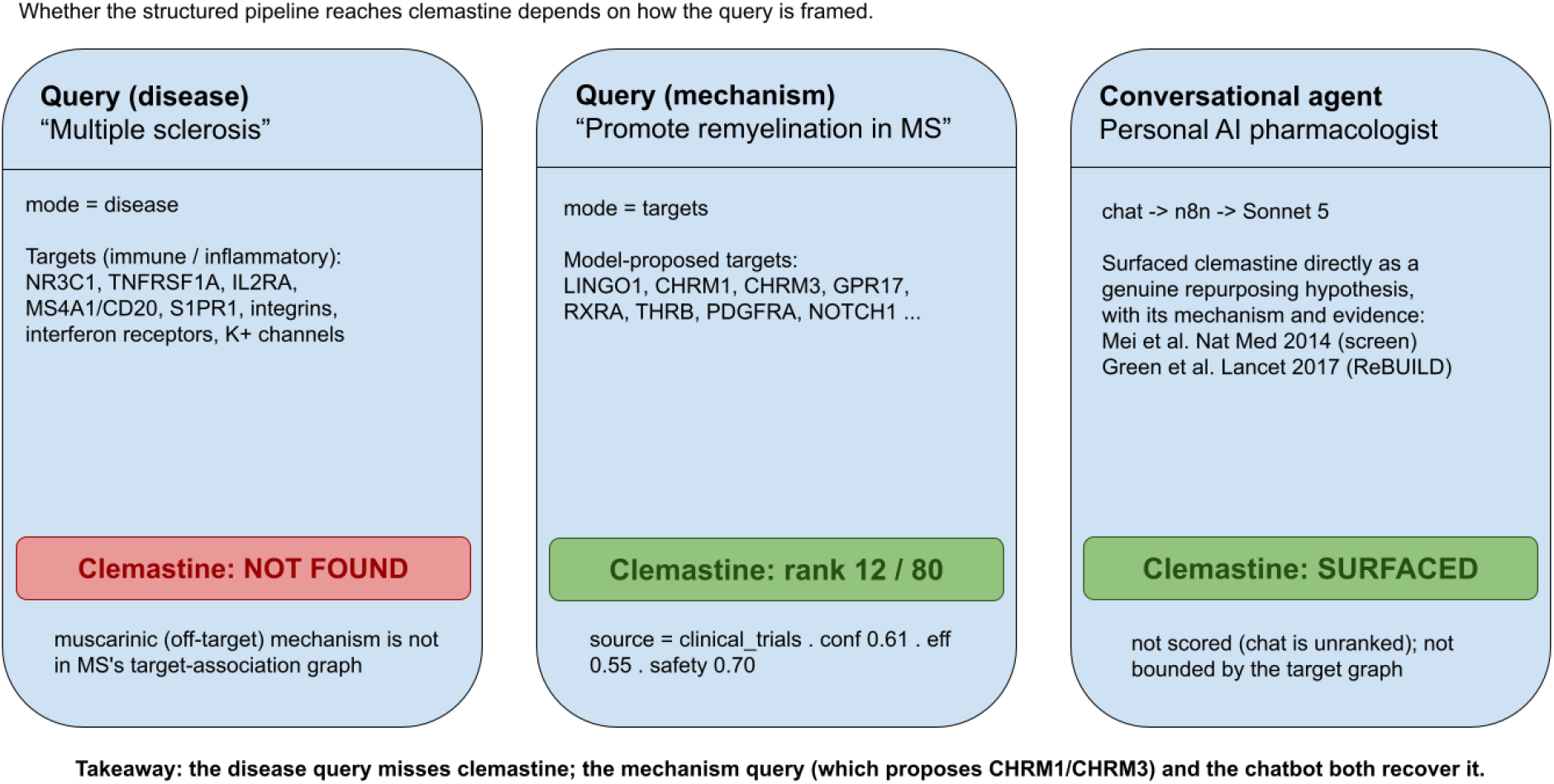
Clemastine in the RePurpose results for multiple sclerosis.

## Supplementary material

*Supplementary File S1*. Exact query strings and conversational-assistant prompts, summaries of the RePurpose reports for each case-study query, and the tunable-ranking analysis for clemastine.

*Supplementary File S2*. How RePurpose Works: a product and technical overview of the tool, including the pipeline, scoring formulas, data sources, and API.

## Notes

https://www.repurposepharma.com

https://github.com/nayagamnitin-hub/RePurpose-Pharma

### Competing Interest Statement

The authors have declared no competing interest.

