## Supplemental File 1 for "RePurpose: An LLM-Assisted, Source-Traceable Drug Repurposing Tool with Retrospective Validation on Clemastine in Multiple Sclerosis"

### Supplementary File S1 — Complete query reports

*Companion to the case study "clemastine in multiple sclerosis."*

| | |
|---|---|
| **Tool / version** | RePurpose, code commit 81f67e4 (2026-10-10) |
| **Model** | Claude Sonnet 5 (anthropic/claude-sonnet-5) via OpenRouter |
| **Runs** | October 2026; one run per structured query; proposer temperature ≈ 0.2 |

These are the archived model-on runs used in the main text. Deterministic steps (interpretation, target resolution, database/trial retrieval, score recalibration) reproduce exactly; the model-proposal step samples at temperature ≈ 0.2, so the AI-proposed subset and clemastine's exact rank in the goal-mode queries are run-specific (see §3.6). For the multiple-sclerosis disease query, the established list and resolved targets are reproduced in full; the 80 repurposing candidates are summarised by source with the two model-proposed entries and the high-confidence candidates listed in full.

#### S1.1 — Exact queries and prompts

**Structured queries.** Disease mode: "multiple sclerosis"; "demyelinating disease". Goal/targets mode: "promote remyelination in multiple sclerosis"; "remyelination / oligodendrocyte precursor differentiation".

##### Conversational-assistant prompts and responses

###### Prompt 1

> Give me THREE genuine drug-repurposing hypotheses. STRICT rules: each must be a drug ALREADY FDA-approved (or long-established) for one disease, now plausibly repurposed for a COMPLETELY DIFFERENT disease area. Do NOT include drugs whose original/primary development was for the proposed new indication (that is not repurposing). For each give: drug name, its CURRENT approved indication, the proposed NEW indication, the mechanistic link, and whether any published studies support it. Favor less-obvious, interesting candidates over textbook examples like aspirin or metformin.

###### Response 1

> The agent proposed three hypotheses: (1) Auranofin (approved: rheumatoid arthritis -> proposed: oncology/antiparasitic; thioredoxin-reductase inhibition; early Phase I/II + preclinical). (2) Itraconazole (approved: antifungal -> proposed: oncology; Hedgehog/Smoothened inhibition, anti-angiogenesis; Phase II signals). (3) Disulfiram (approved: alcohol use disorder -> proposed: glioblastoma/TNBC with copper; copper-dependent NPL4/p97 inhibition; Skrott et al., Nature 2017 + Phase I/II).

###### Prompt 2

> Those three are solid but fairly well-known. Give me TWO MORE that are genuinely surprising and lesser-known (avoid metformin, aspirin, sildenafil, thalidomide, minoxidil, propranolol, and the three you just named). Each must be a drug APPROVED for one disease, repurposed for a clearly DIFFERENT disease area, with a real mechanistic link and published

> evidence. Then, across ALL five, name the SINGLE strongest and most unique repurposing hypothesis and defend it with specific published evidence.

###### Response 2

> The agent proposed two more: (4) Clemastine (approved: OTC H1 antihistamine, allergy -> proposed: remyelination in MS; OFF-TARGET muscarinic M1/M3 antagonism drives OPC differentiation, not its H1 action; Mei et al., Nature Medicine 2014 phenotypic screen; Green et al., Lancet 2017 ReBUILD trial, positive VEP-latency biomarker). (5) Bexarotene (approved: cutaneous T-cell lymphoma -> proposed: Alzheimer's; RXR -> ApoE/ABCA1 amyloid clearance; Cramer et al., Science 2012; FAILED to replicate in humans). Across all five, the agent named CLEMASTINE the single strongest and most unique, on the grounds that it is the only one with a positive randomized placebo-controlled human trial on an objective biomarker, has a doubly-surprising off-target mechanism, and is category-defining (almost no approved MS drug promotes remyelination).

#### S1.2 — Structured report: “multiple sclerosis” (disease mode)

**Interpretation:** mode = disease; disease_terms = multiple sclerosis, relapsing-remitting multiple sclerosis, demyelinating disease. **Resolved disease:** EFO_0803536 (“multiple sclerosis symptom measurement”); 6,347 associated targets, top 30 (score ≥ 0.1) kept. **Overview note:** “Multiple sclerosis has no known cure...”

**Clemastine: NOT FOUND (muscarinic receptors CHRM1/CHRM3 are not among the resolved targets).**

**Counts:** 31 established; 80 repurposing (database 57, clinical_trials 20, AI 2, established 1); opposite-effect 0.

###### Resolved targets (top 30)

| # | Symbol | Assoc. score |
|---|---|---|
| 1 | NR3C1 | 0.6906 |
| 2 | TNFRSF1A | 0.682 |
| 3 | IL2RA | 0.6803 |
| 4 | KEAP1 | 0.639 |
| 5 | MS4A1 | 0.6248 |
| 6 | S1PR1 | 0.6234 |
| 7 | KCNH8 | 0.6208 |
| 8 | IFNAR1 | 0.6126 |
| 9 | KCNA3 | 0.6115 |
| 10 | IFNAR2 | 0.6113 |
| 11 | ITGA4 | 0.6091 |
| 12 | S1PR2 | 0.6084 |
| 13 | S1PR3 | 0.6075 |
| 14 | ITGB1 | 0.6048 |
| 15 | S1PR5 | 0.6043 |

| | | |
|---|---|---|
| 16 | S1PR4 | 0.6005 |
| 17 | IL7R | 0.5991 |
| 18 | KCNA1 | 0.5986 |
| 19 | KCNQ1 | 0.5975 |
| 20 | KCNC1 | 0.5963 |
| 21 | KCNG4 | 0.5962 |
| 22 | ITGB7 | 0.5961 |
| 23 | DHODH | 0.5953 |
| 24 | KCNQ3 | 0.5927 |
| 25 | KCNH7 | 0.5923 |
| 26 | KCNQ2 | 0.5918 |
| 27 | KCNC3 | 0.5918 |
| 28 | HMGCR | 0.5914 |
| 29 | KCNQ5 | 0.591 |
| 30 | KCNA2 | 0.5907 |

##### Established treatments (31)

| # | Drug | Source | Eff | Safety | Conf | Phase | Via targets |
|---|---|---|---|---|---|---|---|
| 1 | Ocrelizumab | database | 0.85 | 0.65 | 0.85 | 4 | MS4A1 |
| 2 | Natalizumab | database | 0.85 | 0.55 | 0.85 | 4 | ITGA4,ITGB1,ITGB7 |
| 3 | Tysabri | clinical_trials | 0.85 | 0.55 | 0.715 | | |
| 4 | Ozanimod | database | 0.75 | 0.68 | 0.75 | 4 | S1PR1,S1PR5 |
| 5 | Ublituximab | database | 0.75 | 0.65 | 0.75 | 4 | MS4A1 |
| 6 | Siponimod | database | 0.75 | 0.65 | 0.75 | 4 | S1PR1 |
| 7 | Ponesimod | database | 0.75 | 0.65 | 0.75 | 4 | S1PR1 |
| 8 | Methylprednisolone Sod. Succinate | database | 0.75 | 0.6 | 0.75 | 4 | NR3C1 |
| 9 | Fingolimod | database | 0.75 | 0.6 | 0.75 | 4 | S1PR1-5 |
| 10 | Cladribine | established | 0.75 | 0.6 | 0.645 | | |
| 11 | Vumerity | clinical_trials | 0.7 | 0.72 | 0.61 | | |
| 12 | Dimethyl Fumarate | database | 0.7 | 0.7 | 0.7 | 4 | KEAP1 |
| 13 | Monomethyl Fumarate | database | 0.7 | 0.7 | 0.7 | 4 | KEAP1 |
| 14 | Tecfidera | clinical_trials | 0.7 | 0.7 | 0.61 | | |
| 15 | Glatiramer Acetate | clinical_trials | 0.65 | 0.75 | 0.719 | 4 | |
| 16 | PEGINTERFERON BETA-1A | database | 0.65 | 0.7 | 0.65 | 4 | IFNAR1,IFNAR2 |
| 17 | INTERFERON BETA-1A | database | 0.65 | 0.7 | 0.65 | 4 | IFNAR1,IFNAR2 |
| 18 | Teriflunomide | database | 0.65 | 0.7 | 0.65 | 4 | DHODH |
| 19 | Rebif | clinical_trials | 0.65 | 0.7 | 0.719 | 4 | |
| 20 | Interferon Beta Therapy | clinical_trials | 0.65 | 0.7 | 0.575 | | |
| 21 | Interferon Beta | clinical_trials | 0.65 | 0.7 | 0.575 | | |
| 22 | Interferon beta-1b (Betaferon) | clinical_trials | 0.65 | 0.68 | 0.575 | | |
| 23 | Interferon beta-1b (Betaseron) | clinical_trials | 0.65 | 0.68 | 0.575 | | |

| | | | | | | | |
|---|---|---|---|---|---|---|---|
| 24 | Baclofen | established | 0.6 | 0.7 | 0.6 | 4 | |
| 25 | Oral Methylprednisolone | clinical_trials | 0.6 | 0.65 | 0.684 | 4 | |
| 26 | Prednisone | database | 0.6 | 0.6 | 0.6 | 4 | NR3C1 |
| 27 | Corticotropin (ACTH) | clinical_trials | 0.6 | 0.55 | 0.684 | 4 | |
| 28 | Mitoxantrone | established | 0.6 | 0.3 | 0.6 | 4 | |
| 29 | Tizanidine | established | 0.55 | 0.7 | 0.55 | 4 | |
| 30 | Dalfampridine | database | 0.5 | 0.75 | 0.5 | 4 | KCNA1-3,KCNC1/3,KCNG4,KCNH7/8,KCNQ1-3/5 |
| 31 | Fampyra | clinical_trials | 0.5 | 0.75 | 0.47 | | |

**Repurposing candidates — high-confidence subset and the two model-proposed entries**

| # | Drug | Source | Eff | Safety | Conf | Phase | Via targets |
|---|---|---|---|---|---|---|---|
| 1 | Venlafaxine | clinical_trials | 0.88 | 0.7 | 0.88 | 4 | |
| 2 | Fluoxetine | clinical_trials | 0.88 | 0.7 | 0.88 | 4 | |
| 3 | Citalopram | clinical_trials | 0.88 | 0.7 | 0.88 | 4 | |
| 4 | Etrasimod | database | 0.944 | 0.73 | 0.848 | 4 | S1PR1,S1PR4,S1PR5 |
| 5 | Tedisamil | database | 0.943 | 0.73 | 0.846 | 4 | K+ channels |
| 6 | Guanidine Hydrochloride | database | 0.943 | 0.53 | 0.846 | 4 | K+ channels |
| 7 | Alemtuzumab | established | 0.85 | 0.5 | 0.823 | 3 | |
| 8 | INTERFERON ALFA-N3 | database | 0.859 | 0.81 | 0.785 | 4 | IFNAR1,IFNAR2 |
| 9 | Vedolizumab | database | 0.858 | 0.52 | 0.783 | 4 | ITGA4,ITGB7 |
| 10 | Fluprednisolone | database | 0.787 | 0.73 | 0.758 | 4 | NR3C1 |
| 11 | Remibrutinib | clinical_trials | 0.76 | 0.6 | 0.76 | 3 | |
| 12 | MD1003 (high-dose biotin) | clinical_trials | 0.76 | 0.6 | 0.76 | 3 | |
| 13 | Daclizumab | ai | 0.75 | 0.3 | 0.75 | | IL2RA |
| 14 | Omaveloxolone | ai | 0.7 | 0.75 | 0.7 | | KEAP1 |

**Model-proposed (AI) repurposing candidates, with rationale**

- **Omaveloxolone** — Approved Nrf2 activator (Friedreich's ataxia) acting through KEAP1; mechanistically parallel to dimethyl fumarate, supporting antioxidant/anti-inflammatory repurposing in MS.
- **Daclizumab** — Previously approved for relapsing MS via CD25 (IL2RA) blockade; directly validates IL2RA as an MS target despite later withdrawal for safety.

#### S1.3 — Structured report: “demyelinating disease” (disease mode)

**Interpretation:** mode = disease. **Clemastine: NOT FOUND**; no muscarinic (CHRM) targets resolved. Same pattern as the multiple-sclerosis disease query.

#### S1.4 — Structured reports: goal/targets mode (clemastine recovered)

##### Query (goal/targets mode): “promote remyelination in multiple sclerosis”

**Model-proposed targets:** LINGO1, CHRM1, CHRM3, RXRA, THRB, PDGFRA, NOTCH1, GPR17, HDAC1, ESR1, IGF1, CNTF, BDNF. **Muscarinic:** CHRM1, CHRM3.

**Clemastine: FOUND in repurposing at rank 12/80 (source clinical_trials; eff 0.55, safety 0.7, conf 0.613).**

**Model-proposed (AI) candidates this run:** Sobetirome (GC-1), PIPE-307, UAB30, Crenigacestat, Nirogacestat (PF-03084014), VK2809, Clobetasol, Telenzepine, Temelimab (GNbAC1), Elezanumab, Miconazole, High-dose Biotin (MD1003).

##### Query (goal/targets mode): “remyelination / oligodendrocyte precursor differentiation”

**Model-proposed targets:** LINGO1, CHRM1, CHRM3, HRH3, GPR17, RXRA, THRB, PPARG, NOTCH1, BMP4, MTOR, PDGFRA. **Muscarinic:** CHRM1, CHRM3.

**Clemastine: FOUND in repurposing at rank 76/80 (source clinical_trials; eff 0.45, safety 0.65, conf 0.543).**

**Model-proposed (AI) candidates this run:** Sobetirome, VK2809, Telenzepine, Crenigacestat.

#### S1.5 — Tunable-ranking (slider) analysis for clemastine

Clemastine entered the “promote remyelination” list as a clinical_trials candidate (max_phase 3, confidence 0.613). novelty = 0.5·(1−phase/4) + 0.5·source_weight; display = α·novelty + (1−α)·confidence.

| α | Clemastine rank (/80) | Display score |
|---|---|---|
| 0.0 | 15 | 0.613 |
| 0.25 | 9 | 0.566 |
| 0.5 | 12 | 0.519 |
| 0.75 | 21 | 0.472 |
| 1.0 | 21 | 0.425 |

The rank is non-monotonic (peaks at α ≈ 0.25): a phase-3 trial drug has only moderate novelty, so maximum novelty weighting demotes it.
