## Supplemental File 2 for "RePurpose: An LLM-Assisted, Source-Traceable Drug Repurposing Tool with Retrospective Validation on Clemastine in Multiple Sclerosis"

### RePurpose

How RePurpose Works - Product & Technical Overview

*Nitin Nayagam · Sirpi Internship Project · 2026*

*Live at repurposepharma.com*

#### What's new in this version

Four things have been added since the last version of this document:

- **Tunable sort for repurposing candidates -** a slider from well-established to under-explored, with a Sort button, so a 80-100-candidate list can be re-weighted toward novelty without ever discarding confidence.
- **Live pipeline status while a search runs -** the loading line now rotates through what the pipeline is actually doing, instead of a generic “Analyzing...”.
- **A built-from-scratch Snake game -** fades in if a search is still running after about four seconds, so there is something to do besides watch a spinner.
- **Clickable protein targets -** every protein chip now opens a popup with the protein's function and a link to UniProt.

#### The idea

Most drugs do more than one thing. Minoxidil was developed to treat severe high blood pressure, until doctors noticed patients growing hair and it became a leading hair-loss treatment. Sildenafil was studied for chest pain before its side effect made it famous as Viagra. These second lives are common, and finding them by hand means reading across gene databases, drug registries, trial listings and PubMed at once.

RePurpose does that traversal automatically. You give it a condition, a target, or a plain-English goal; it works out which proteins drive that biology, which direction each one needs to be pushed, then pulls every drug that acts on those proteins and sorts them by whether they push the right way. It is a hypothesis-generation and literature-exploration tool, not a source of medical advice.

#### What the tool does

You give it a condition (Alzheimer's disease), a target (EGFR), or a plain goal (how to lose body weight). The result is deliberately split in two.

- **Established treatments -** the approved, real-world options.
- **Repurposing and in-development candidates -** the interesting half: drugs in active clinical trials for related conditions, plus mechanistically plausible leads the model proposes from the confirmed target biology.

Anything acting in the wrong direction is moved into an opposite-effect list rather than quietly dropped, so you can always see what was excluded and why.

#### While you wait

A search is not instant. The loading state shows what is actually happening rather than a generic spinner: the status line shimmers and rotates through the real pipeline stages as they run, the same rotating-status pattern used in the Personal AI Pharmacologist's own loading animation.

```
Interpreting your query... → Mapping the biology on Open Targets... → Finding
drugs that act on those targets... → Searching active clinical trials... →
```

```
Proposing novel repurposing leads... → Scoring effectiveness and safety... →
Reading PubMed evidence... → Writing your plain-English summary...
```

If a search is still running after about four seconds, a small Snake game fades in beneath the status line. It is built from scratch on an HTML canvas, with no external libraries and no borrowed assets (Google's dinosaur game was deliberately not cloned, since that is Google's IP; Snake is a generic classic that is safe to implement from scratch), and styled in the site's own warm palette: clay-orange food, charcoal snake. It plays with arrow keys or WASD on desktop and swipe gestures on mobile, keeps a live score, and offers a Restart button.

The moment the report is ready, or the search errors out, the game and its timers and key listeners are torn down cleanly so nothing bleeds into the results page, and the footer stays put.

One game was chosen deliberately over several, to keep the page light and focused: Snake is instantly familiar, so there is nothing new to learn while waiting. A second option, such as a simple 2048 or memory/tile game with a small toggle to choose between them, is an easy addition later if it's worth doing.

#### What comes back

A report has four parts, each traceable to a source database:

- **Targets explored** - the proteins the condition runs through, each with an Open Targets association score and a plain-language function pulled from UniProt.
- **Established treatments** - approved options, ranked by effectiveness then safety, with mechanism of action, clinical stage and known indications.
- **Repurposing and in-development candidates** - ranked by confidence, each marked with where it came from: a structured database, an active clinical trial (with its NCT numbers), or an AI-proposed mechanistic lead.
- **Opposite effect** - drugs that hit the right targets the wrong way, kept visible for transparency.

Because a goal-style search can return 80 to 100 repurposing candidates, that list carries a tunable sort control rather than one fixed order: a slider running from Well-established to Under-explored, with a Sort button beside it. At its default position the list matches the original confidence-based order exactly. Moving the slider and clicking Sort blends each candidate's confidence score with a novelty measure built from clinical phase and evidence source (structured database, active trial, or AI-proposed lead), so pushing toward Under-explored surfaces earlier-stage and AI-proposed leads that a confidence-only sort tends to bury behind “show more,” without ever discarding confidence altogether. The exact blend is given later, in How the scores are built.

Every drug card carries clickable badges - Approved, Experimental, Black-box warning, Most effective, Safest. Clicking a badge asks the model to justify that specific claim and pulls real PubMed studies for it, so anything on a card can be chased back to literature. Each card also offers View studies (a live PubMed search) and Ask AI (a question answered from those study snippets, with PMIDs cited).

Protein targets are interactive in the same spirit. Every protein chip, in Proteins targeted at the top of a goal-style report and in Targets explored at the bottom, is clickable, marked with a small info icon and a hover highlight. Clicking one opens a popup with the protein's full name and function; FGFR3, for example, resolves to “Fibroblast growth factor receptor 3, a tyrosine-protein kinase that acts as a cell-surface receptor for fibroblast growth factors and regulates cell proliferation...,” plus a link to the full entry on UniProt. It closes with the close button, a click outside the popup, or the Esc key.

A dedicated /api/protein endpoint powers this: one cached UniProt lookup per protein, with no LLM call involved, so it stays fast and does not draw on the OpenRouter budget.

#### The Personal AI Pharmacologist

A conversational research assistant focused on pharmacology and drug repurposing. Talk to it like a person, get a plan built for your goal, or hand it a study to break down. It stays on-topic: pharmacology and research only.

It reasons mechanistically, reads studies you hand it, and builds plans around a goal. There are two entry points, and they behave differently on purpose:

- **Personal AI Pharmacologist** (under the search bar) - the full consult page. Always a clean slate: it knows nothing about a search you ran earlier.
- **Talk to AI** (top right) - a docked chat you can use while reading results. This one is screen-aware: while results are in front of you it knows what you searched, so “which of these is furthest along” resolves correctly.

Context is only helpful when it matches what you are looking at, so leaving the results clears it, and a New chat control wipes the transcript, the history and the session id the assistant keys its memory on.

##### What you can ask it

- **Talk naturally** - a real back-and-forth about drugs, mechanisms and research in plain language, down to the pharmacokinetics: receptors, pathways, onset, half-life, CYP metabolism, and why a dose behaves nonlinearly.
- **Build your plan** - ask it to design a specific plan, for example lose weight and gain muscle, or regrow hair, grounded in the biology rather than a listicle.
- **Analyze studies** - paste a study (PMID, PMCID, DOI or PubMed URL) and get the key takeaways, plus how the findings could translate into future or repurposing uses. The backend fetches the actual text from NCBI, so factual claims are based on that fetched text; repurposing potential is reasoned separately and labelled as a hypothesis.

##### The interaction rule

Suggesting a drug without its interaction profile is the failure mode that actually hurts someone: propranolol for stimulant-driven tachycardia reads as sensible until you notice modafinil inhibits CYP2C19 and slows propranolol's clearance. Before proposing anything to add, combine or switch, the assistant is therefore required to state the clinically significant interactions with whatever you have said you are taking, naming the mechanism (which enzyme and in which direction, transporters, additive sympathomimetic, QT or serotonergic load), placing the warning beside the suggestion rather than in a footnote, and saying plainly when something needs a clinician.

#### How it is wired

Chat posts to /api/chat, which keeps the assistant's webhook URL server-side. The backend assembles the message - on-screen context when applicable, the last few turns for pronoun resolution, any fetched study text, a style directive and the interaction rule - and sends it to an n8n agent workflow, running Claude Sonnet 5, that holds the topic guard and long-term memory. Greetings and “what can you do” are answered locally so the topic guard never rejects a normal opener, and if n8n is unreachable the request falls back to the app's own OpenRouter key (Claude Sonnet 5) with a leaner prompt, so the chat keeps working.

#### The pipeline, step by step

One search runs this sequence. The order matters: each stage narrows or enriches what the next one sees.

**1. Interpret the query, with direction.** The model turns free text into targets and the direction each should move. “Grow taller” means boost the GH/IGF axis but suppress FGFR3 and oestrogen-driven growth-plate fusion. Encoding direction is what lets later stages keep agonists and drop antagonists rather than listing anything that merely touches a relevant protein.
*llm/interpret.py · QueryInterpretation*

**2. Resolve the biology.** Disease names resolve to an EFO id and then to ranked associated targets; gene symbols resolve straight to Ensembl ids. Targets below an association score of 0.1 are dropped and the top 30 are scanned.
*Open Targets GraphQL · EFO · Ensembl*

**3. Describe the proteins.** The leading targets are enriched with their human protein function and names, which makes the mechanism explanations readable rather than a list of symbols.
*UniProt REST*

**4. Collect every drug that acts on them.** For each target, up to 30, the known drugs with mechanism of action, action type, clinical phase, approved indications, warnings and adverse-event signals, all drawn from Open Targets' drug data.
*Open Targets*

**5. Filter by direction.** Each drug's action type is reduced to a sign: agonist, activator and opener count as positive; inhibitor, antagonist, blocker and degrader as negative, and checked against the direction the goal wants. Wrong-way drugs move to the opposite-effect list.
*services/pipeline.py · goal_fit*

**6. Add what is actually being trialled.** A ClinicalTrials.gov v2 search over the goal's conditions surfaces agents no static database lists yet. Because these are being trialled for the goal, they are inherently goal-appropriate, and each keeps its NCT ids.
*ClinicalTrials.gov v2 · NCT ids*

**7. Propose beyond the databases.** Grounded on the confirmed targets and the real drugs that hit them, the model proposes more selective or next-generation agents the registries do not link yet. These are flagged prospective and AI-proposed so they read as hypotheses.
*llm/propose.py · source: ai*

**8. Collapse duplicates, then reassess.** Brand names, salts and formulations collapse to one entry per active ingredient. A final clinical-assessment pass recalibrates effectiveness and safety, drops off-goal entries, and adds well-known drugs the databases missed. A deterministic guardrail then runs regardless of what the model said: nothing unapproved and below phase 4 may sit in “established.”
*llm/assess.py · guardrail*

**9. Write the narrative.** Scores and badges are assigned, and the model writes the summary and a note on whether the condition has a cure or only management. Studies are fetched per drug on demand rather than up front, which keeps the initial search fast.
*NCBI E-utilities · BioC*

Finished reports are cached per query, so repeat searches return instantly and deterministically; a refresh=true parameter forces a recompute.

#### How the scores are built

Three numbers drive the ordering and the badges. They are heuristics, and worth understanding before you trust the sort order.

#### Effectiveness and confidence

For a goal-style query, where directional fit is the strongest available signal:

```
effectiveness = 0.50·goal_fit + 0.20·(phase / 4) + 0.30·establishment
confidence = 0.60·effectiveness + 0.40·goal_fit
```

For a disease-style query, where target coverage carries more weight:

```
effectiveness = 0.60·(phase / 4) + 0.25·target_coverage + 0.15·target_relevance
confidence = 0.70·effectiveness + 0.30·target_relevance
```

#### Safety, and why it is not a report count

Counting FAERS adverse-event reports is wrong, because report volume mostly tracks how widely a drug is prescribed: metformin would look dangerous and an obscure phase-1 compound pristine. Safety here instead scores how well characterised and how benign the profile is, then clamps the result to a floor and ceiling:

| Factor | Effect | Reasoning |
|---|---|---|
| Approved (phase 4) | base 0.78 | Established, characterised profile |
| Phase 3 / 2 / 1 | 0.60 / 0.45 / 0.32 | Progressively less known |
| Preclinical | base 0.20 | Risks largely unknown, so low rather than high |
| Black-box warning | -0.30 | Serious documented harm |
| Withdrawn | -0.55 | Pulled from market |
| Each additional warning | -0.04 | Accumulated caution flags |
| Oncology / cytotoxic class | -0.20 | Inherently harsher class |
| Small molecule / biologic | -0.05 / +0.03 | Biologics tend to be better tolerated |
| Breadth of adverse-event types | up to -0.18 | Distinct signals, capped, not raw volume |

These formulas are not the final word on a live search. When a model is configured, a late assessment pass overwrites effectiveness and safety with its own medical judgment and recomputes confidence, so the formulas above act as the deterministic base and as the fallback used when no model is available.

#### Tuning the sort: well-established vs. under-explored

The repurposing-candidates list adds one more layer on top of confidence: a slider that blends confidence with how novel a candidate is, so the same underlying scores can be viewed two ways.

```
novelty = 0.5·(1 - phase/4) + 0.5·source_weight
display_score = alpha·novelty + (1 - alpha)·confidence
```

source_weight is 1.0 for an AI-proposed lead, 0.6 for a candidate that comes from an active trial only, and 0.0 for one already listed in a structured database. alpha is the slider's position: 0 (well-established) reproduces the original ranking exactly, 1 (under-explored) weights fully toward novelty. The list only re-sorts when the user releases the slider and clicks Sort, computed client-side from the candidates already on the page, so moving the slider itself never disturbs the visible order until they confirm it.

### Technical details

#### Data sources

Every source is free and public. Only NCBI takes an optional key, which raises the rate limit from 3 to 10 requests per second.

| Source | Role in the pipeline | Endpoint |
|---|---|---|
| Open Targets Platform | Structured backbone: disease → ranked targets, target → drugs with mechanism, action type, clinical phase, warnings and adverse events | api.platform.opentargets.org/api/v4/graphql |
| ClinicalTrials.gov v2 | Drugs actually under investigation for the goal's conditions, with NCT ids | clinicaltrials.gov/api/v2 |
| UniProt | Protein function and names for each target | rest.uniprot.org/uniprotkb |
| NCBI E-utilities | Topic or drug → PubMed ids | eutils.ncbi.nlm.nih.gov/entrez/eutils |
| NCBI BioC | PMID → abstract, PMCID → open-access full text, as clean JSON | ncbi.nlm.nih.gov/research/bionlp/RESTful |
| PMC ID Converter | PMID ↔ PMCID, so a reference always resolves to fetchable text | ncbi.nlm.nih.gov/pmc/utils/idconv/v1.0 |
| NCBI Citation Exporter | Formatted citations (RIS / MEDLINE) | api.ncbi.nlm.nih.gov/lit/ctxp/v1 |
| Wikipedia REST | Plain-language drug description and photo for the profile card | en.wikipedia.org/api/rest_v1/page/summary |
| PubChem PUG-REST | 2D chemical structure image | pubchem.ncbi.nlm.nih.gov/rest/pug |

#### Stack

**Backend** - Python 3.12, FastAPI, uvicorn, Pydantic v2 with pydantic-settings for env-driven config. HTTP through httpx, deliberately synchronous: FastAPI runs sync handlers in a threadpool, so the client code stays readable without blocking the loop.

**Frontend** - hand-written HTML, CSS and vanilla JavaScript served straight from the app. No framework, no build step, no bundler.

**Structure** - one module per external API under app/clients/, LLM work isolated in app/llm/, orchestration in app/services/pipeline.py, and the response contract in app/models/schemas.py.

**Legacy UI** - a Streamlit app remains in the repo from earlier development, but it is not part of the deployed site.

#### Models

The app depends only on an LLMProvider interface with a single complete() method, so the model behind it is a one-line swap. The provider layer ships classes for OpenRouter, Groq, Gemini, and Ollama, plus a Mock fallback, chosen by the LLM_PROVIDER setting with key validation. It is currently set to OpenRouter, running anthropic/claude-sonnet-5 for both the analysis pipeline and the conversational assistant. If no valid key is configured it falls back to the Mock provider, so the pipeline still returns the structured report with the AI narrative layers quietly disabled rather than inventing text.

##### HTTP API

| Route | Access | Does |
|---|---|---|
| GET /api/health | open | Health check used by the host |
| GET /api/config | open | Public Turnstile site key and enabled flag |
| POST /api/verify | open | Validates a Turnstile token, sets the human cookie |
| GET /api/repurpose?q= | gated | Runs the whole pipeline, returns the cached report |
| GET /api/studies?drug=&topic= | gated | PubMed studies for one drug and topic |
| POST /api/ask | gated | Question about a drug, answered from study snippets with PMIDs |
| POST /api/explain | gated | Badge or score explanation |
| POST /api/evidence | gated | “Learn more” behind a specific badge, with matching studies |
| POST /api/drug | gated | Drug profile card: description and images |
| GET /api/protein?symbol= | gated | Protein name and function for a target chip; one cached UniProt lookup, no LLM call |
| POST /api/chat | gated | The AI pharmacologist, proxied to n8n with fallback |

Gated routes sit behind a Cloudflare Turnstile bot-check: after a one-time verification the server sets a signed cookie that those routes require. Health, config, and verify stay open.

##### Deployment

The live site is at repurposepharma.com, a custom domain routed through Cloudflare to the Render service. RePurpose runs on Render as a Python 3.12 web service rooted at backend/, built from requirements-web.txt and started with uvicorn app.main:app; the health check is /api/health. Configuration and secrets, the OpenRouter key, the n8n webhook URL, and the Cloudflare Turnstile keys, live in environment variables set in the Render dashboard and are never committed, and the service auto-deploys on every push to main. Locally it runs the same command on port 8000, with interactive API docs at /docs.

#### What it is not

Read this before trusting an answer. RePurpose is a research and education tool for generating and exploring hypotheses. It is not medical advice, not a diagnosis, and not a claim that any listed drug works for the goal you typed.

- **Prospective means unproven.** Trial-backed and AI-proposed candidates are labelled as such precisely because nobody has shown they work for that goal. A high confidence score means the mechanism lines up, not that the outcome is established.
- **The scores are heuristics.** Clinical phase is a crude stand-in for effectiveness, and the safety score models how well characterised a drug is, not a measured risk.
- **The interaction rule is instruction, not screening.** The assistant is required to raise interactions, which catches the common documented ones, but it is not a validated interaction database. Anything you would actually take belongs in front of a pharmacist or physician first.
- **The model can be wrong.** Structured facts trace back to source databases; the narrative layers are generated, and generated text can be confidently mistaken.
